# Phylogenomic analyses of the Austral podocarps (*Podocarpus*: Podocarpaceae) reveals unlikely hybrid ancestry of a New Zealand species

**DOI:** 10.64898/2026.02.25.708080

**Authors:** Raees Khan, Ed Biffin, John Conran, Robert Hill, Kor-jent van Dijk, Michelle Waycott

## Abstract

Hybridisation is ubiquitous amongst plants and has important evolutionary consequences ranging from the collapse of distinct lineages through to the generation of new species. Here, we develop a phylogenetic hypothesis for the Austral podocarps (*Podocarpus*), a monophyletic group of six species distributed in Tasmania, mainland Australia, New Zealand and New Caledonia, and identify a putative hybrid lineage. Using a targeted capture approach to generate DNA sequence data, we find discordance between nuclear and plastid derived phylogenetic estimates and in particular, the relationships of the New Zealand species *Podocarpus nivalis* and Australian *P. lawrencei* are significantly discordant. Species network analyses largely resolve this incongruence and indicate that *P. nivalis* is a hybrid lineage, with *P. laetus* (New Zealand) and *P. lawrencei* as parents. We hypothesise that *P. nivalis* has arisen following trans-Tasman dispersal of *P. lawrencei,* and shows eco-geographic divergence from *P. laetus,* which could facilitate reproductive isolation. We suggest that introgression from *P. laetus* to colonising *P. lawrencei* could significantly reduce founder effects while cold tolerance inherited from *P. lawrencei* has enabled *P. nivalis* to occupy alpine environments.

Our findings highlight the importance of reticulate evolution in Southern Hemisphere conifers and demonstrate the value of phylogenomic network approaches for resolving recent and complex radiations.

## Introduction

Hybridisation (i.e., reproduction between species) is a widespread and common phenomenon amongst plants (Whitney et al. 2010) and has diverse and complex evolutionary outcomes ranging from hybrid unviability to the generation of new species (Abbott et al. 2013). Novel lineages may form when hybridisation is accompanied by genome doubling, with the resulting allopolyploid having a level of intrinsic reproductive isolation from parental types (Rieseberg and Willis, 2007). In contrast, homoploid hybrid speciation occurs without a change in ploidy and is considered less common than allopolyploid speciation. This, in part, reflects the absence of intrinsic reproductive isolation in newly formed homoploid hybrids which, initially rare, are prone to demographic swamping by parental species and the elimination of hybrid phenotypes (Soltis and Soltis, 2009). Nevertheless, several convincing cases of homoploid hybrid speciation have been documented (Long and Rieseberg, 2024) including amongst conifers (Mao et al. 2011; Sun et al., 2014).

Ecogeographic separation from parental species is generally observed among putative homoploid hybrid species (Gross and Rieseberg, 2005; Abbott et al. 2010) and is required to achieve reproductive isolation from the parental species (Buerkle et al. 2000). However, the frequency of homoploid hybrid species is poorly known in part reflecting the diverse array of outcomes arising from hybridisation other than speciation (e.g., introgression; Long and Rieseberg, 2024). Schumer et al. (2014) suggest a set of criteria for the recognition of homoploid hybrid species including that reproductive isolation is a direct consequence of hybridisation, which has been considered overly restrictive by others (Nieto Feliner et al. 2017).

A history of hybridisation is suggested by discordance between gene trees, and in particular, conflicting estimates from nuclear and plastid derived markers (cytonuclear discordance) are often interpreted as evidence for gene flow.

Chloroplast capture, the process of inter-lineage transfer of the chloroplast following hybridisation and repeated backcrossing, provides one explanation for cytonuclear conflict (but see Larson et al. 2024). In angiosperms, the chloroplast is usually inherited maternally and because there is a negative correlation between rates of intraspecific gene flow and interspecific introgression (Currat et al. 2008; Petit and Excoffier, 2009), seed-dispersed plastids are prone to introgression across morphological species boundaries (Rieseberg and Soltis, 1991). In contrast, chloroplast inheritance in conifers is usually paternal (Owens and Wilson, 2003; Neale and Wheeler, 2019), which confers high rates of intraspecific gene flow.

Consequently, chloroplast variation among conifers is more likely to be species-specific, and this is supported by empirical studies (e.g., Du et al. 2009; Zhou et al. 2017.).

Cytonuclear discordance can arise due to introgression affecting either the cytoplasmic or nuclear genome compartments, and due to factors other than hybridisation including methodological errors (e.g., gene tree estimation error, undetected paralogy) or biological processes such as incomplete lineage sorting (ILS). It is therefore important to determine whether incongruence reflects alternative underlying histories for genes or genomes or can be ascribed to other factors (Folk et al. 2017; Garcia et al. 2017; Larson et al. 2024). In the era of genome scale data and datasets comprising hundreds to thousands of nuclear loci, the distribution of gene trees (or sites) relative to a species tree estimate can be used to distinguish ILS and gene flow (Martin et al. 2015; Blischak et al. 2018). Similarly, approaches that explicitly model gene flow in the presence of ILS use multilocus data to infer explicit phylogenetic networks (e.g., Solís-Lemus and Ané, 2016; Solís-Lemus et al. 2017). While full likelihood and Bayesian approaches are computationally demanding, approaches based upon maximum pseudo-likelihood scale to large datasets and are compatible with goodness-of-fit tests to determine model fit to the data (Solís-Lemus and Ané, 2016; Cai and Ané, 2021).

The New Zealand biota has been considered variously as a vicariant Gondwanan relict or derived entirely by long distance dispersal (Pole et al. 1994; Winkworth et al. 2002) following the Oligocene marine transgression event and associated mass extinction (Cooper and Cooper, 1995). While recent research suggests a more nuanced view (e.g., Wallis and Jorge, 2018), there is little debate regarding the significance of long-distance dispersal in the assembly of the modern New Zealand biota and in particular, trans-Tasman dispersals from Australia to New Zealand are relatively frequent (Jordan, 2001; Winkworth et al. 2002; Nge et al. 2021).

Nevertheless, extreme long-distance dispersal is improbable (Nathan, 2006) and there are several barriers to the establishment of a viable population following such an event (Wu et al. 2023). Newly arrived immigrants are vulnerable to Allee effects, and this is particularly true for self-incompatible species, such as dioecious plants (Baker, 1955; Elam et al. 2007; Levin et al. 2009). However, when an alien species encounters a more abundant interfertile resident there is an enhanced probability of hybridisation (Abbott and Reisberg, 2001; Beddows and Rose, 2018; Mitchell et al. 2019), which can have both short (e.g., heterosis) and longer term (e.g., adaptive variation) benefits for colonists due to increased genetic variation (Rius and Darling, 2014; Pfennig et al. 2016).

*Podocarpus* is the largest genus within the predominantly South Hemisphere distributed conifer family Podocarpaceae, and is divided into two subgenera: subgenus *Podocarpus,* which includes predominantly high latitude temperate Southern Hemisphere taxa; and subgenus *Foliolatus,* which is more diverse in tropical Asia and Malesia (de Laubenfels, 1985). Within subgenus *Podocarpus*, a group of seven species with small, narrow flattened leaves and predominantly high latitude distributions was recognised by de Laubenfels (1985) as section *Australis*. Of these, four species from New Zealand: *P. acutifolius, P. laetus, P. nivalis, P. totara*) and one species each from Australia (*P. lawrencei*) and New Caledonia (*P. gnidioides*) form a monophyletic group in recent molecular phylogenies (e.g., Biffin et al. 2012; Knopf *et al*., 2012; Quiroga *et al*., 2016) (hereafter, Austral podocarps; Figure 1). *Podocarpus nubigenus* is also included in sect. *Australis* by de Laubenfels (1985), but a close relationship to the remaining species is not supported by molecular data (e.g., Stull et al. 2021; Adaïmé et al, 2024) and will not be considered further. As with the family more generally, the Austral podocarps are dioecious and produce a fleshy female cone suggesting endozoochory (Khan and Hill, 2022). To date, phylogenetic estimates for the group have been based upon limited sampling of taxa and of genetic data, and interspecific relationships are poorly understood. The New Zealand species often occur in close geographic proximity and the widespread occurrence of inter-specific hybrids complicates the assignment of individuals to species (Wardle, 1972; Webby et al. 1987; Marshall et al. 2015).

**Figure 1:**
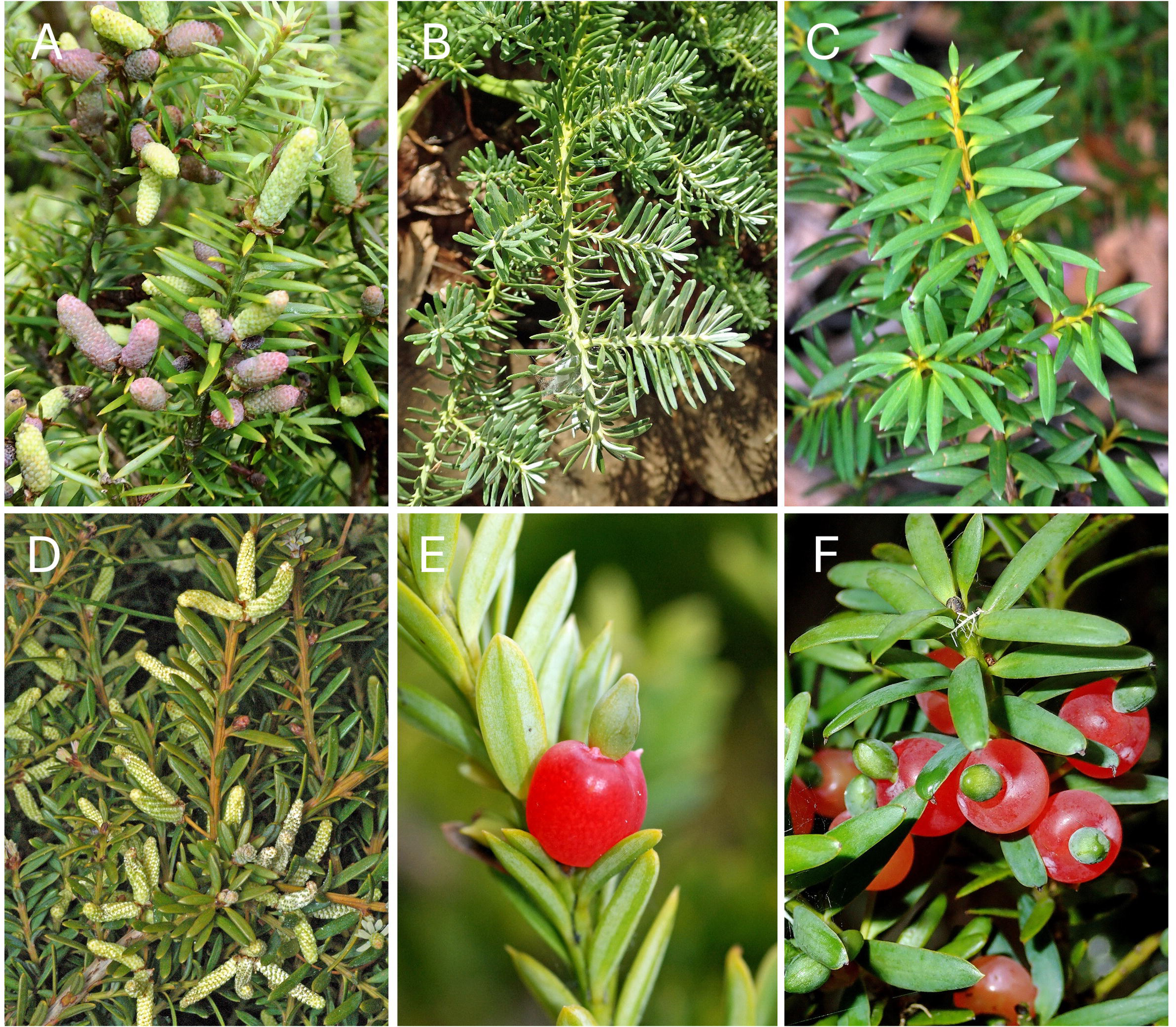
Examples of the Austral podocarps. A. *Podocarpus acutifolius* (pollen cones), Dunedin Botanic Gardens; B. *P. gnidioides*, Institut Agronomique Néo-Calédonien, St Louis, New Caledonia; C. *P. laetus* Mt Lofty Botanic Gardens W932295; D. *P. lawrencei* (pollen cones), Mt Lofty Botanic Gardens G888806; E. *P. nivalis* (seed cones), Dunedin Botanic Gardens; F. *P. totora* (seed cones), Omeru Reserve, Kaukapakapa, New Zealand. All photographs taken by J.G. Conran.

In the present study, we use hybrid capture and high throughput sequencing to develop a phylogenomic hypothesis for the Austral podocarps, comprising the Australian, New Zealand and New Caledonian representatives of de Laubenfels sect. *Australis.* We compare datasets generated from both nuclear and plastid genome compartments and provide evidence for a hybrid origin of *P. nivalis* in New Zealand following trans-Tasman dispersal.

## Materials and methods

### DNA extraction and sequencing

Samples included a mixture of herbarium, and fresh leaf material collected into silica gel (Table 1). DNA extraction and library preparation were performed at the Advanced DNA, Identification and Foresnsics Facility (ADIFF; University of Adelaide). DNA was extracted with the Qiagen Plant Mini kit, normalised to 2 ng/uL and was sheared using a Diagenode Bioruptor® Pico sonicator to fragment lengths of c. 400–600 bp. DNA libraries were constructed using a JetSeq Flex DNA Library preparation kit (Bioline). Each sample was assigned a unique combination of two synthetic barcodes, each of 8 bp length, which were annealed at each end of the DNA fragments.

**Table 1:** Samples included in this study. Sample codes for individuals included in the ‘reduced’ dataset for network and HyDe analyses are italicised and bold sample codes indicate individuals not included in the plastid dataset. All herbarium specimens were sourced from the Dame Ella Campbell Herbarium (MPN), Massey University, New Zealand.

| Taxon | Sample code | Collection code/voucher number | Population | Location |
| --- | --- | --- | --- | --- |
| *Podocarpus lawrencei* Hook.f. | TNK25 | NK25 | Australia-NSW | Mount Jagungal |
|  | TMK02 | MK02 | Australia-NSW | Thredbo River |
|  | TMP3 | MP3 | Australia-VIC | Mount Pinnibar |
|  | VEB2 | EB2 | Australia-VIC | Goonmirk Rocks |
|  | VRK1 | RK1 | Australia-VIC | Mount Baw Baw |
|  | VRK5 | RK5 | Australia-VIC | Mount Baw Baw |
|  | VRK14 | RK14 | Australia-VIC | Mount McKay |
|  | TMF06 | MF06 | Australia-Tasmania | Mount Field |
|  | TPL03 | PL03 | Australia-Tasmania | Pine Lake |
|  | TNE08 | NE08 | Australia-Tasmania | Ben Lomond Plateau |
|  | TCPO8 | CP08 | Australia-Tasmania | Cradle Mountain |
|  | TCPO9 | CP09 | Australia-Tasmania | Cradle Mountain |
|  | TME04 | ME04 | Australia-Tasmania | Mersey river |
|  | TMR05 | MR05 | Australia-Tasmania | Mount Read |
| *Podocarpus nivalis* Hook. | PNMS1 | PNMS1 | New Zealand-South Island | Gertrude Saddle |
|  | PNMS2 | PNMS2 | New Zealand-South Island | Gertrude Saddle |
|  | PNMS3 | PNMS3 | New Zealand-South Island | Gertrude Saddle |
|  | PNMS4 | PNMS4 | New Zealand-South Island | Gertrude Saddle |
|  | **PNMS5** | PNMS5 | New Zealand-South Island | Gertrude Saddle |
|  | MPN1 | MPN27453 | New Zealand-North Island | Volcanic Plateau |
|  | **MPN2** | MPN43974 | New Zealand-North Island | Waiouru Army Training Group Land |
|  | MPN4 | MPN48619 | New Zealand-South Island | Temple Basin, Canterbury |
|  | MPN5 | MPN38282 | New Zealand-South Island | Red Hills Ridge |
|  | PN 59 | - | cultivated | Mount Lofty Botanical Garden |
| *Podocarpus totara* G.Ben ex D.Don. | PTN1 | MPN51493 | New Zealand-North Island | Palmerston North |
|  | **PTS2** | MPN45404 | New Zealand-South Island | Bankview Station "Long Bush Gully" |
|  | PTN4 | MPN51310 | North Island | Wedde Wood |
|  | PTS5 | MPN39626 | cultivated | Dunedin Waipouri Forest |
|  | PT62 | - | cultivated | Mount Lofty Botanical Garden |
| *Podocarpus acutifolius* Kirk | PAN2 | MPN41818 | New Zealand-North Island | Taupō |
|  | PAN3 | - | cultivated | Mount Lofty Botanical Garden |
|  | PAN4 | - | cultivated | Mount Lofty Botanical Garden |
| *Podocarpus laetus* Hooibr. ex Endl. | PLN2 | MPN43969 | New Zealand-North Island | Rangipo |
|  | *PLS4* | PLS4 | New Zealand-South Island | Gertrude Saddle |
|  | *PLS5* | PLS5 | New Zealand-South Island | Gertrude Saddle |
| *Podocarpus gnidioides* Carrière | *PNN1* | - | cultivated | Mount Lofty Botanical Garden |
|  | *PNN2* | - | cultivated | Mount Lofty Botanical Garden |
| *Podocarpus salignus* D.Don | OUT | - | cultivated | Mount Lofty Botanical Garden |

We used a conifer-specific bait set targeting 100 low-copy nuclear loci (Khan et al. 2024) and a chloroplast bait set targeting c. 30 plastid coding regions that were designed on angiosperms (Waycott et al. 2021) using the MYBaits target enrichment system (MYcroarray, Ann Arbor, Michigan) to the enrich samples for the selected loci. Hybrid capture was performed following the manufacturer’s instructions (https://arborbiosci.com/genomics/targeted-sequencing/mybaits/) with the hybridisation step using an annealing temperature of 65 ◦C and incubation for 24 hr. Post capture PCR was performed on the half-built libraries to fuse the remaining sequencing adapters with Illumina P5 and P7 indexes. Libraries were pooled in equimolar concentrations and sent for Illumina paired-end sequencing (2×150) on a lane of a HiSeqX Ten at the Garvan Institute for Medical Research in Sydney.

### Bioinformatics processing

High-throughput paired-end reads were imported into CLC Genomics Workbench (ver. 20.0.2, QIAGEN, see https://digitalinsights.qiagen.com/) for demultiplexing and trimming using a quality score limit of 0.05 (Phred score ∼13). Reads for each individual were randomly sampled to 2.5 million paired-end reads.

We used CAPTUS v1.1.1 (Ortiz et al. 2023; Raza et al, 2023) to *de novo* assemble the sequencing reads for each individual, extract nuclear and plastid target regions from the assemblies and generate multiple sequence alignments (MSAs) for each extracted gene region. For the assembly step the preset ‘CAPSKIM’ settings (see https://edgardomortiz.github.io/captus.docs/assembly/assemble/options/index.html) for Megahit (v.1.2.9; Li et al. 2015) were used, with otherwise default settings. For the extraction step the sequences originally used to design the nuclear probe set (REMcon; Khan et al. 2024) were used as targets with minimum identity and minimum score set at 80% and 0.5, respectively. To extract the plastid regions, we generated a reference set comprising coding sequences from *Podocarpus lambertii, P. latifolius, P. milanjianus* and *P. totara* (GenBank accession numbers NC023805, NC042224, MT019686, NC020361, respectively) and used minimum identity and minimum score values of 90% and 0.5, respectively. For the alignment step we used MAFFT (v. v7.526; Katoh et al. 2002) with the *mafft-auto* algorithm and *informed* paralog filtering with a *tolerance* of 1 to generate ‘genes-flanked’ (coding sequences+introns+flanking regions) MSAs for the nuclear and plastid data.

For the nuclear data, we used the pre-paralog filtering alignments output from CAPTUS as input to the Python package ParaGone v1.1.3 (https://github.com/chrisjackson-pellicle/ParaGone; Jackson et al. 2023). ParaGone implements the methods described by Yang and Smith (2014), which generate quality-controlled alignments and rooted homolog gene trees as input to tree-based orthology inference. We used MAFFT and TAPER (Zhang et al. 2021) to generate cleaned MSAs from the paralog files and Fasttree (v2.1; Price et al. 2010) to generate gene trees, with *Podocarpus salignus* designated as an internal outgroup. Treeshrink v.1.3.9 (Mai and Mirarab, 2018) was used to detect and remove outlier long branches from the gene trees using a *q* value of 0.2, and the monophyletic outgroups algorithm was used to generate ortholog alignments.

### Phylogenetic analyses

Maximum likelihood phylogenies were estimated in IQ-Tree 2 (Minh et al. 2020) for each of the nuclear and plastid data. For the nuclear phylogeny, we used a concatenated matrix of 92 loci output from ParaGone including 38 individuals (37 ingroup samples). For the plastid analyses, the concatenated alignment included 29 chloroplast loci including 35 individuals (34 ingroup samples) (Table 1). For both datasets, we used ModelFinder (MFP + MERGE, see http://www.iqtree.org/ModelFinder/) to estimate the best nucleotide substitution model for each locus and merge loci with similar models into a single partition (Kalyaanamoorthy et al. 2017). To reduce computational burden, only the top 10% of partition merging schemes was considered (Lanfear, 2014). The best fitting model and partitioning scheme was then used in subsequent analyses with 1000 ultrafast bootstrap replicates (UFBS; Minh et al. 2013) to assess branch support.

For the nuclear dataset, we used IQ-Tree 2 to estimate gene trees for each locus. For these analyses, ModelFinder was used to estimate the best nucleotide substitution model for each partition and we used 1000 UFBS replicates to measure branch support. This set of gene trees was used to estimate a species tree under the multi-species coalescent model, which is statistically consistent in the presence of incomplete lineage sorting (ILS) (Mirarab et al. 2014). For these analyses, we used the weighted Astral algorithm (wASTRAL; Zhang and Mirarab, 2022) as implemented in Accurate Species Tree Estimator (ASTER, ver. 1.4.2.3, see https://github.com/chaoszhang/ASTER). This algorithm down weights poorly supported quartets and long branches in gene trees, thereby reducing the impact of phylogenetic uncertainty in species tree estimates (Zhang and Mirarab, 2022). For these analyses, the bootstrap support value ‘preset’ was used (see https://github.com/chaoszhang/ASTER/ tutorial/astral-hybrid.md) to define maximum and minimum values for bootstrap support, and branch support on the species tree was measured using local posterior probabilities (Sayyari & Mirarab, 2016).

Disagreement between the nuclear and plastid phylogenies was visualised using the tanglegram function of the dendextend R package (Galili, 2015; https://github.com/talgalili/dendextend).

### Concordance analyses

We used PhyParts (Smith et al. 2015) to explore patterns of conflict among the nuclear gene trees and the species tree estimate. For a given target tree (i.e., the species tree), PhyParts summarises the number of genes that are concordant with, conflict with, or are uninformative with respect to each node in the target tree topology. Here, we used the ASTRAL tree as the target and mapped the nuclear gene trees against this topology. Additionally, we mapped the nuclear gene trees against the plastid topology to assess whether there is support from the latter in instances of cyto-nuclear discord (e.g., Fu et al. 2023). The PhyParts output was visualized using a Python script (phypartspiecharts.py by M. Johnson, available at: https://github.com/mossmatters/phyloscripts/tree/master/phypartspiecharts).

Quartet Sampling (QS) (Pease et al. 2018) provides branch support measures that attempt to quantify underlying variation in the data, given that for large phylogenomic datasets, high levels of support are expected using conventional measures such as the bootstrap (Lanfear and Hahn, 2024). QS generates three metrics for internal nodes on the target topology: quartet concordance (QC) describes the proportion of concordant versus both discordant quartet topologies; quartet differential (QD) describes skewness of discordant topologies; and quartet informativeness (QI) quantifies the proportion of informative replicates (Pease et al., 2018). We performed a maximum of 1000 QS replicates with the log-likelihood threshold cutoff was set to 2 and the –*calc_qdstats* flag activated to assess the significance of QD statistics using the Chi-square test, with significant values considered indicative of gene flow (Pease et al. 2018; Cai et al, 2021). The R script plot_QC_ggtree.R (by ShuiyinLIU, available at: https://github.com/-ShuiyinLIU/QS_visualization) was used to visualised the QS results.

### Gene tree simulations

We followed the approach of Folk et al. (2017), Garcia et al. (2017) and others to assess whether a hypothesis of ILS is sufficient to explain topological discordance between plastid and nuclear phylogenies for *Podocarpus.* Using the species tree estimate from ASTRAL as a guide, we first simulated 1000 gene trees under a coalescent model, with branch lengths scaled by a factor of four to approximate those expected for the plastid among a dioecious diploid lineage (Small et al. 2004). We used DendroPy v. 5.0.1(Sukumaran et al. 2010) and the wrapper script simulate_gene_trees.py (by R. Folk, available at; https://github.com/ryanafolk/tree_utilities) to perform the simulations. The simulated trees were then mapped onto the empirical plastid phylogeny using Phyparts and visualised using the phypartspiecharts.py script. The null hypothesis is that cyto-nuclear discordance is explained by ILS. In this case, the empirical plastid topology should be recovered with reasonable frequency among the simulated plastid topologies. The alternative is that inferred plastid relationships are unexpected under ILS alone, indicating that gene flow provides a reasonable additional explanation for plastid-nuclear conflicts. Under this scenario, empirical plastid relationships should be absent or found in very low frequency (e.g., ≤ 1%; Thureborn et al. 2024) among the simulated gene trees.

### Assessment of gene flow

We estimated a network topology for *Podocarpus* using the Species Networks applying Quartets (SNaQ) method as implemented in the Julia package PhyloNetworks v1.0.0 (Solís-Lemus and Ané, 2016; Solís-Lemus et al. 2017). This approach infers species network topologies based upon quartet concordance factors derived from gene trees under maximum pseudolikelihood in the presence of gene flow and ILS (Solís-Lemus and Ané, 2016). To reduce computation burden, we analysed a dataset including 18 ingroup samples including representatives of species groups/clades recovered in the full data set analyses (Table 1). Gene trees and an ASTRAL species tree were generated for the reduced sample as detailed above. The gene trees were used to estimate quartet concordance factors, and the ASTRAL topology was used as input to estimate a topology under ILS alone (*hmax*=0). The resulting topology was used for a network search adding one hybridisation edge, and network searches with up to 5 hybridisation edges (*hmax*=5) were conducted, each using the optimal network from the previous step. We conducted 10 independent runs of SNaQ to infer the optimal network for each *hmax* value. The fit of networks was assessed using the log pseudolikelihood profile with *hmax* (Solís-Lemus and Ané, 2016). We also used the Julia package QuartetNetworkGoodnessFit (Cai and Ané, 2021) to compare the distributions of gene tree quartet topologies with those expected for a candidate network. Topologies with higher *p*-values are considered more likely, corresponding to a high proportion of well-fitting quartet trees. These tests were conducted for each *hmax* value using 1000 simulations of *Z* scores and optimised branch lengths.

In addition we used HyDe 0.4.3 (Blischak et al., 2018) a coalescent method that can detect gene flow in the presence of ILS using site frequency patterns among triplets of ingroup taxa relative to an outgroup taxon (Kubatko and Chifman, 2019), similar to ABBA-BABA tests. HyDe estimates the amount of admixture (γ), which ranges from 0-1, with γ values of c. 0.5 indicative of a 50:50 hybrid while values closer to 0 and 1 suggest low levels of asymmetric introgression. Using the concatenated 92 nuclear gene data set with *Podocarpus gnidioides* as the outgroup, we tested all possible four-taxon combinations using the reduced (18 samples; Table 1) dataset, as well as the full (37 samples) data. Populations were defined as in Figure 2. Subsequently, we tested for hybridisation at the individual level within populations that showed significant levels. Significant p-values (*p<*0.05) were corrected using Bonferroni correction, and Z-scores greater than three were taken as strong evidence of hybridization (Blischak et al., 2018).

**Figure 2:**
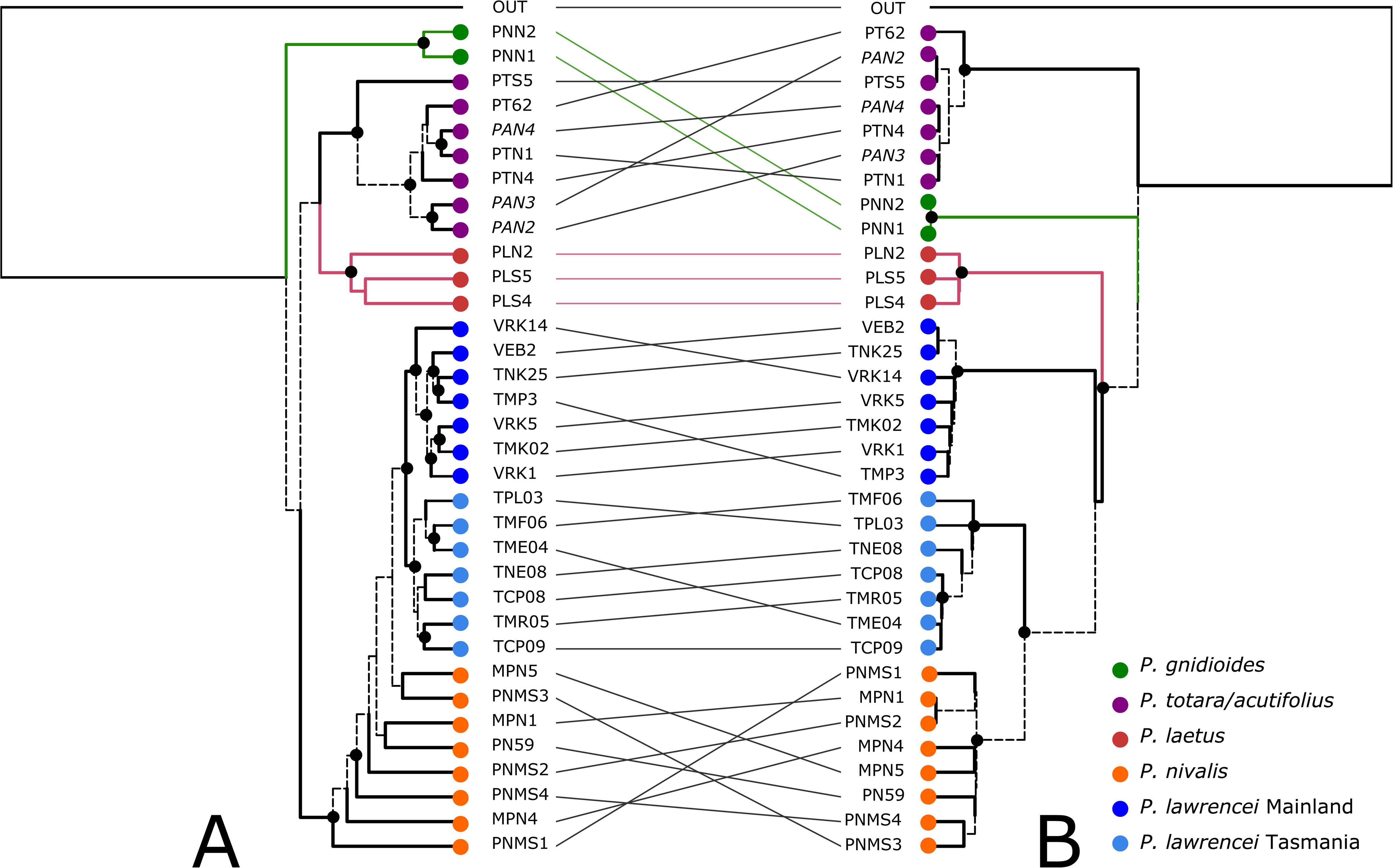
Tanglegram comparing maximum likelihood topologies inferred from (a) concatenated nuclear and (b) plastid datasets (for support values see Figure S1 and S5, respectively). Dashed branches are unique to that tree and coloured branches and connecting lines indicate subgroups common to both trees. Taxon designation is indicated by the coloured circles on terminal branches.

## Results

### Nuclear phylogeny

The concatenated alignment of 92 nuclear gene regions and 38 individuals had a length of c. 140 kbp and approximately 25% missing data (Supplementary Data 1). The average un-gapped length of sequences was c. 105 kbp and the average aligned length of the individual gene regions was c. 1520 bp (range c. 700 to 6000 bp). Species trees estimated from these data using concatenation (IQ-TREE; Figure 2a; Figure S1) and summary species tree approach (ASTRAL; Figure S2) provide similar inferences of relationships among individuals and clades. The monophyly of both *P. gnidioides* and *P. lawrencei* is strongly supported while a clade including *P. totara* and *P. acutifolius* is recovered but neither of these species is resolved as monophyletic. *Podocarpus laetus* is resolved as monophyletic, or weakly paraphyletic in the concatenated versus ASTRAL topologies, respectively, and *P. nivalis* is paraphyletic in both analyses within a strongly supported clade including *P. lawrencei.* Within *P. lawrencei,* a clade comprising mainland individuals (hereafter, mainland clade) is recovered in both analyses while a Tasmanian clade is either monophyletic (IQ-TREE) or weakly paraphyletic (ASTRAL).

Despite the reasonable statistical support for several relationships within the Austral podocarps from both the concatenated and coalescent approach, we found signals of strong discordance among individual nuclear genes. With respect to the Phyparts analyses, many bipartitions in the species tree were rarely, if ever, recovered among individual genes. For instance, most of the splits within the *P. nivalis–P. lawrencei* clade were recovered at frequencies of zero to <10% among gene tree topologies, with the majority recovering low frequency alternative topologies (Figure S3).

Similarly, quartet sampling indicates low support (QC values close to 0) for the many relationships in the species tree including several negative values. These include nodes that show significant QD values (Chi-square test P-value <0.05) indicating that one discordant topology is favoured over the other (Pease et al. 2016) (Figure S4).

### Plastid phylogeny

The plastid dataset included 29 gene regions and 36 individuals with an aligned length of c. 51 kbp (average un-gapped length of 47 kbp) and approximately 7% missing values (Supplementary Data 2). The average aligned length per locus was 1700 bp, ranging from c. 600-5000 bp. The topology inferred from the concatenated alignment using IQ-TREE is shown in Figure 2b (Figure S5). We found strong support for species groups including *P. gnidioides, P. laetus* and *P. nivalis.* As with the nuclear topologies, we recovered a well-supported clade including *P. acutifolius* and *P. totara* while intraspecific sampling indicates non-monophyly of both species. A key difference between the nuclear and plastid topologies is the resolution of *P. lawrencei,* which in the latter includes *P. nivalis* as sister to Tasmania *P. lawrencei,* while the mainland clade is sister to these. Also contrasting the nuclear topologies, *P. laetus* is resolved as sister to the *P. nivalis–P. lawrencei* clade, rather than with *P. totara–P. acutifolius* (Figure 2a).

### Coalescent Simulations

Using the rescaled nuclear species tree as a guide, we simulated a distribution of organellar gene trees under a coalescent model (Supplementary Data 3). When these were mapped to the empirical plastid topology (Figure 3, Supplementary Data 3), we found that some discordant relationships were recovered with reasonably high frequency among simulated gene trees, exceeding the threshold to reject ILS as a sufficient explanation for cyto-nuclear differences. For instance, a monophyletic *P. laetus* was found in c. 15% of simulated topologies, the placement of *P. laetus* as sister to *P. nivalis* and *P. lawrencei* was found in c. 3% of simulations and the monophyly of Tasmanian *P. lawrencei* had a frequency of c. 2.5%. On the other hand, several plastid relationships were never recovered, or found at very low frequency, among the simulated gene trees. These include the monophyly of *P. nivalis* (c. 0.8% of simulations), the monophyly of *P. nivalis* and Tasmanian *P. lawrencei* (not found in any simulations) and relationships among several samples within the *P. totara–P. acutifolius* clade.

**Figure 3:**
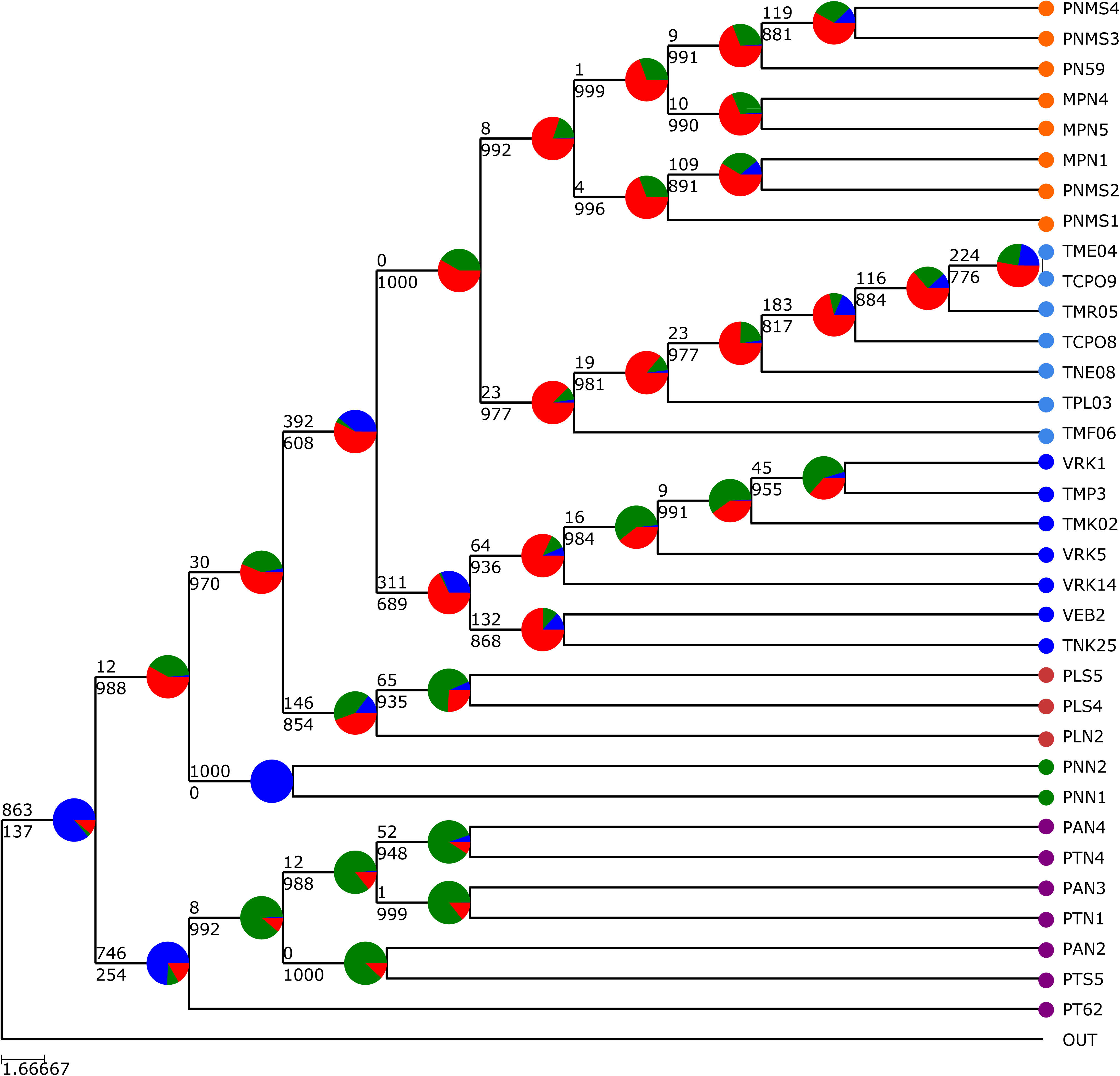
Empirical plastid topology (Figure 1b) showing the distribution of 1000 plastid topologies simulated under a coalescent model and mapped using PhyParts. Numbers of concordant and discordant simulated gene trees are show above and below each branch, respectively. The pie charts represent the proportion of concordant (blue) and discordant (main alternative, green; all other alternatives, red) topologies for that branch. Taxon designation as in Figure 1.

### Hybridisation Assessment

Network inferences using SNaQ with 18 terminals found up to four hybridisation edges with both the goodness of fit tests (Table 2) and log pseudolikelihood profile plots (Figure 4c) favouring a network topology over a fully bifurcating tree. Each of the inferred networks suggests a hybrid origin of *P. nivalis* with parentage including *P. laetus* (PLS5) and the ancestor of the *P. lawrencei* mainland clade, with inheritance probabilities of c. 49% and 51%, respectively (Figure 4a-b). Additional hybridisation edges were recovered within *P. nivalis,* the *P. lawrencei* mainland clade and within the *P. totara–P. acutifolius* clade up to an *hmax* value of 3 (Figure 4b) and no further reticulations were added for *hmax* values of 4 and 5. The goodness of fit tests were inconclusive regarding the best network topology, with *hmax* values from 1-5 providing an adequate fit (*P*-value > 0.05; Table 2). The plot of log pseudolikelihood and *hmax* suggests little improvement in model likelihood above *hmax=*2 (Figure 4c).

**Figure 4:**
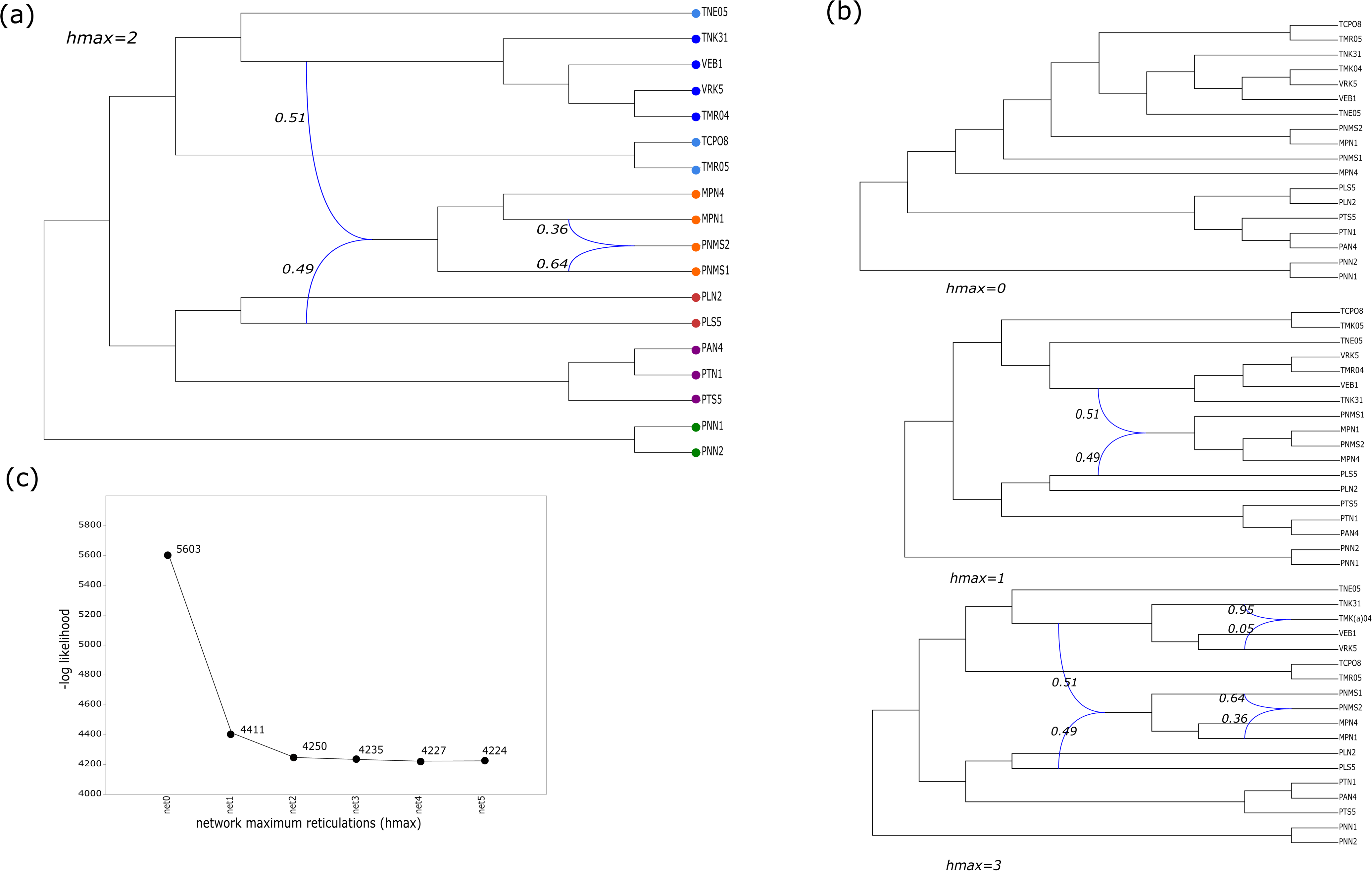
Phylogenetic network analysis for the reduced (18 terminal) dataset using SNaQ. Species networks inferred allowing (a) 2, (b) 0, 1 and 3 reticulations. The pseudo-loglikelihood scores (c) show little improvement for *hmax* values greater than 2. Blue curved branches indicate the possible hybridisation event and inheritance probabilities are shown adjacent to that branch. Taxon designation as in Figure 1.

**Table 2:** Results of the goodness of fit tests, showing the model tested, the negative pseudo log-likelihood value and significance value are indicated for each *hmax* value. Topologies with higher *p*-values are considered more likely, corresponding to a high proportion of well-fitting quartet trees.

|  | model | pseudo loglikelihood |
| --- | --- | --- |
| net0 | ILS | -5603.45 |
| net1 | ILS+1 hybrid edge | -4411.51 |
| net2 | ILS+2 hybrid edge | -4249.764 |
| net3 | ILS+3 hybrid edge | -4234.986 |
| net4 | ILS+4 hybrid edge | -4226.7 |
| net5 | ILS+5 hybrid edge | -4224.43 |

For the reduced (18 ingroup) data, 2 out of 60 hypotheses tested using HyDe showed significant evidence of hybridisation (after Bonferroni correction, *Z* scores > c. 3.6). Both identified *P. nivalis* as the hybrid taxon with either the *P. lawrencei* mainland (0.31< γ < 0.69) or Tasmanian clades (0.33< γ < 0.67) and *P. laetus* identified as parental species. For the full data set, a single significant hypothesis again identified a hybrid ancestry for *P. nivalis* between Tasmanian *P. lawrencei* and *P. laetus* (0.44< γ < 0.56). At the individual level, 3 out of 4 *P. nivalis* samples in the 18 taxon data had a *Z* score > 3.5, providing strong evidence for hybrid ancestry, while for the 38 sample data, 4 out of 10 *P. nivalis* individuals had *Z* scores exceeding 3.5 (Table 3).

**Table 3:** HyDe results with significant γ values. For each set of parental taxa (P1, P2), a γ value significantly different from 0 indicates a signal of hybridisation in the ‘Hybrid’ taxon.

**HyDe results full dataset**
| P1 | Hybrid | P2 | Zscore | Pvalue | γ |
| --- | --- | --- | --- | --- | --- |
| *P.lawrencei* (Tas) | PNMS3 | *P. laetus* | 4.7668462 | 9.37E-07 | 0.51 |
| *P.lawrencei* (Tas) | PNMS4 | *P. laetus* | 4.53588626 | 2.87E-06 | 0.6 |
| *P.lawrencei* (Tas) | MPN2 | *P. laetus* | 3.94864449 | 3.93E-05 | 0.52 |
| *P.lawrencei* (Tas) | MPN5 | *P. laetus* | 3.78003929 | 7.84E-05 | 0.57 |

**HyDe results 18 terminals**
| P1 | Hybrid | P2 | Zscore | Pvalue | γ |
| --- | --- | --- | --- | --- | --- |
| *P.lawrencei* (Mainland) | PNMS1 | *P. laetus* | 3.90357197 | 4.74E-05 | 0.69 |
| *P.lawrencei* (Mainland) | PNMS2 | *P. laetus* | 3.76554708 | 8.31E-05 | 0.7 |
| *P.lawrencei* (Mainland) | MPN1 | *P. laetus* | 4.22785245 | 1.18E-05 | 0.7 |
| *P.lawrencei* (Tas) | PNMS1 | *P. laetus* | 4.04416926 | 2.63E-05 | 0.63 |
| *P.lawrencei* (Tas) | PNMS2 | *P. laetus* | 3.7808132 | 7.82E-05 | 0.66 |
| *P.lawrencei* (Tas) | MPN1 | *P. laetus* | 3.99492111 | 3.24E-05 | 0.72 |

## Discussion

Our phylogenetic analyses of the Austral podocarps revealed strongly conflicting topologies from nuclear versus plastid genes, and particularly the resolution of *P. nivalis.* While the nuclear topologies resolve *P. nivalis* as a paraphyly including *P. lawrencei* (Figure 2a; Figures S1-2), the plastid data place *P. nivalis* within *P. lawrencei* as sister to the Tasmanian *P. lawrencei* clade (Figure 2b; Figure S5). Provided gene trees are estimated without error (see below) gene tree-species tree discordance can arise due to gene duplication and loss, introgression/hybridisation and ILS. With respect to paralogy, we used an approach that explicitly attempts to infer ortholog groups from homolog gene sets (Yang and Smith 2014; Jackson et al. 2023). We subsequently used coalescent simulations to test support for ILS versus gene flow hypotheses. These analyses suggest that ILS alone cannot account for the observed cyto-nuclear discordance, and specifically, the plastid relationships noted above were never found among plastid topologies simulated on the ASTRAL species tree under ILS (Figure 3). While not ruling out a role for ILS, these findings suggest that explanations invoking gene flow and/or chloroplast introgression are also required to explain cytonuclear discordance (Folk et al. 2017).

Hybridisation has been widely reported amongst conifers (Neale and Wheeler, 2019) and both natural and artificial interspecific hybrids have been recorded among the Austral podocarps (Webby et al. 1987; Marshall et al. 2015). Amongst angiosperms, cyto-nuclear discordance is often attributed to introgressive hybridisation leading to chloroplast capture (e.g., Folk et al. 2017; Garcia et al. 2017), which in part is a consequence maternal chloroplast inheritance and lower rates of gene flow relative to biparentally inherited pollen (Currat et al. 2008). For most conifers (including *Podocarpus;* Owens and Wilson, 1999, for *P. totara*) the plastid is inherited paternally, which confers high levels of intraspecific gene flow. Modelling (Currat et al. 2008) and empirical studies (e.g., Zhou et al. 2017) demonstrate a negative correlation between rates of intraspecific gene flow and interspecific introgression suggesting that chloroplast capture is less likely amongst conifers relative to lineages with maternal chloroplast inheritance (Du et al. 2009; Zhou et al. 2017). While several instances of putative chloroplast introgression have been reported for conifers (e.g., Liston et al. 2007; Adams, 2016; Uckele et al. 2021), these generally involve lineages that occur in sympatry or parapatry (Neale and Wheeler, 2019) or allopatric lineages with hypothesised secondary contact, e.g., in historical refugia (Liston et al. 2007). Generally, introgression is asymmetric from resident into ‘invading’ species gene pools (Currat et al. 2008; Neale and Wheeler, 2019). Given that *P. nivalis* and *P. lawrencei* are allopatric across the Tasman Sea (c. 2000 km separation) a chloroplast capture event (i.e., the introgression of *P. lawrencei* chloroplast into *P. nivalis*) seems an unlikely explanation for observed cyto-nuclear discordance.

In addition to cyto-nuclear conflict, we found that nuclear gene trees were also highly discordant with respect to our species trees estimates (Figures S3-4). Factors including gene tree estimation error, ILS and hybridisation/introgression are likely to contribute to discordant gene tree-species tree estimates (Cai et al. 2021) and untangling the relative contribution of these factors can be challenging (Smith et al. 2015; Pease et al. 2016; Rose et al. 2021). In our study, gene tree estimation error and ILS appear to be important sources of incongruence given that many relationships are recent (i.e., intraspecific) and branch lengths are short. Under these conditions, there may be few changes to diagnose clades and limited time between divergence events for lineages to sort (Smith et al. 2015). Nevertheless, we recovered largely congruent topologies using concatenation and coalescent (ASTRAL) frameworks and while the latter relies on individual gene tree estimates to infer the species tree, the concatenation approach does not. Both approaches may be inaccurate in the presence gene flow. A phylogenetic network approach that simultaneously models ILS and interspecific gene flow (Degnan and Rosenberg 2009; Solis-Lemus et al. 2016;) probably provides the best available estimate of the evolutionary history of the *Podocarpus* sect. *Australis* clade (Figure 4a).

Several recent studies have found that the phylogenetic placement of hybrid lineages is often intermediate to the parents and may form ladder-like grades reflecting the genomic contribution of parental species (e.g., Dolinay et al. 2021; Pyron et al. 2022). Interestingly, while the nuclear species tree estimates resolve a paraphyletic *P. nivalis* (Figure 2a, Figures S1-2), the best fitting network topologies resolve this species as monophyletic (Figure 4). The major tree (i.e., the tree derived by deleting minor hybrid edges; Figure S6) extracted from the network places *P. nivalis* within *P. lawrencei*, providing partial resolution of incongruence between the plastid and nuclear data inferred in the absence of gene flow estimates. A consistent interpretation of the plastid topology is that *P. nivalis* represents a hybrid lineage with *P. lawrencei* as the pollen parent (Figure 2b, Figure S3).

The PhyloNetworks and HyDe analyses identified *P. nivalis* as a hybrid with parental contributions from the New Zealand species *P. laetus* and Australian *P. lawrencei*.

Both tests suggest a near equal contribution of genetic material from each parent (γ ∼ 0.5; Figure 4, Table 3), which has been considered a signature of hybrid speciation (Jiggins et al. 2008; Schumer et al. 2014). Spatial and/or ecological separation from parental species is generally observed among putative homoploid hybrid species (Gross and Rieseberg, 2005; Abbott et al. 2010), and in the absence of intrinsic barriers to gene flow, is required to achieve reproductive isolation from the parental species (Buerkle et al. 2000). *Podocarpus nivalis* is spatially isolated from one parent and occurs in a distinct habitat relative to the other. Both *P. lawrencei* and *P. nivalis* have the highest level of freezing resistance recorded for the genus (Sakai and Wardle, 1978; Sakai et al. 1981), which is reflected in core ranges that are centred on alpine and subalpine environments. In contrast, *P. laetus* has lower freezing tolerance (Sakai and Wardle, 1978) with a range extending from lowland to lower subalpine environments (Wardle, 1972) and is largely absent from above the treeline.

While genetic evidence of hybridisation is a prerequisite for hybrid speciation hypotheses, it may be difficult to distinguish several among alternative scenarios without causal evidence linking reproductive isolation to hybridisation (e.g., introgression; Schumer et al. 2014; Long and Rieseberg, 2024; but see Nieto Feliner et al. 2017). In the present case, it seems likely that cold tolerance evolved first in *P. lawrencei,* and therefore, cannot be considered a consequence of hybridisation for *P. nivalis.* Rather, this would suggest that a *P. lawrencei* ancestor first dispersed to New Zealand and has been introgressed by *P. laetus.* An outcome of long-distance dispersal beyond a species core range is that founding populations comprising one or few propagules are subject to strong stochastic effects, which limit establishment success (Blackburn et al. 2015), and these may be particularly pronounced for obligate-outcrossing species such as dioecious plants (Elam et al. 2007). However, where a coloniser encounters an interfertile resident species and abundances are skewed towards the latter, there is an increased likelihood of hybridisation (Abbott and Reisberg, 2001; Beddows and Rose, 2018; Mitchell et al. 2019). Hybridisation can result in increased genetic variation in the founding lineage, which can ameliorate the negative effects of bottlenecks by increasing heterozygosity (i.e., heterosis; Rius and Darling, 2014; Pfennig et al., 2016). In the longer term, increased genetic variance in admixed populations may facilitate adaptation to novel environments (Endelman and Mallet, 2021). In the context of our evidence, hybridisation between colonising *P. lawrencei* and a resident *P. laetus* may have facilitated establishment of the former, while the occupation of high elevation environments by *P. nivalis* suggests a selective advantage conferred by cold tolerant *P. lawrencei* parentage. Eco-geographic barriers are thought to play a key role in preserving the distinctiveness of sympatric and parapatric lineages in insular floras (Crawford and Archibald, 2017; Reatini and Vision, 2024) and may help to maintain species boundaries, despite evidence of ongoing gene flow between *P. nivalis* and *P. laetus* (Webby et al. 1987).

## Conclusions and further work

We found patterns of incongruence between nuclear and plastid phylogenies for the Austral podocarps, which can be explained at least in part by incorporating hybridisation into evolutionary hypotheses. Specifically, our findings indicate that *P. nivalis* has arisen following a trans-Tasman dispersal event(s) involving *P. lawrencei* and subsequent hybridisation with *P. laetus.* While *P. nivalis* shows a genetic signal that is consistent with a homoploid hybrid origin, we cannot rule out alternative hypotheses (e.g., introgression of *P. laetus* genes into *P. lawrencei*) given that cold tolerance in *P. nivalis* was likely inherited from *P. lawrencei.* Distinguishing among these hypotheses is difficult and may involve establishing whether genes derived from both parental species have contributed to reproductive isolation (e.g., Ren et al. 2012) and requires adequate population and genetic sampling.

Many New Zealand lineages have putative Australian origins, including multiple dispersals within a single lineage (e.g., Nge et al. 2021) as well as conspecific trans-Tasman disjunctions (Jordan, 2001). In this light, further instances of secondary contact leading to heterospecific mating seem plausible with the outcome depending on the (lineage specific) degree of divergence among gene pools (Cerca et al. 2023; Reatani and Vision, 2024). While extreme long-distance dispersal events are unlikely, small population effects may also present significant barriers to the establishment of newly arrived immigrant taxa. In the present case, gene flow from a resident to colonising species appears to have facilitated establishment of a novel lineage and may be critical given that *P. lawrencei* is dioecious. Hybridisation has been extensively documented for the New Zealand biota (e.g., Morgan-Richards et al. 2009; Shepherd et al. 2024) although additional studies are needed to assess the frequency and outcomes of secondary contact following long-distance dispersal.

More specifically, New Zealand (in common with other insular floras, see Schlessman et al. 2014; McGlone and Richardson, 2023) has a disproportionately high incidence of dioecious plants relative to the global average despite predictions that obligate outcrossers should be poor colonisers (Baker, 1995) suggesting a more nuanced view of Bakers Law.

## Contributions

All authors contributed to the study conception and design. Material preparation, data collection and analysis were performed by RK, EB and KVD. The first draft of the manuscript was written by EB. RK, JGC, KVD, RM and MW commented on previous versions of the manuscript. All authors read and approved the final manuscript.

---

**Figure S1:**
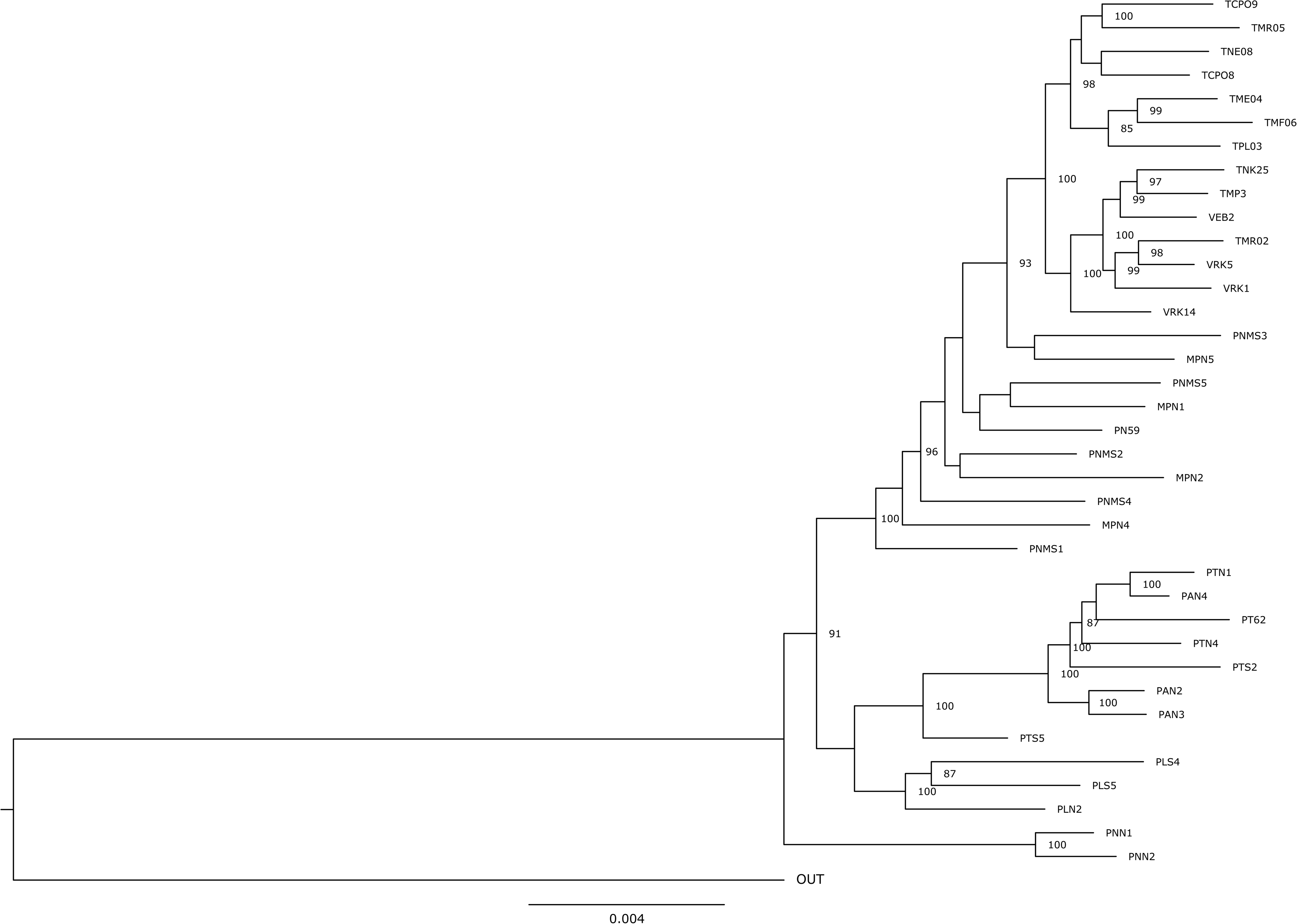
Maximum likelhood topology inferred from the concatenated nuclear dataset using IQ-TREE. Branch lengths are proportional to the number of changes occurring on that branch and support values (UFBS):≥ 80% are shown adjacent to that branch.

**Figure S2:**
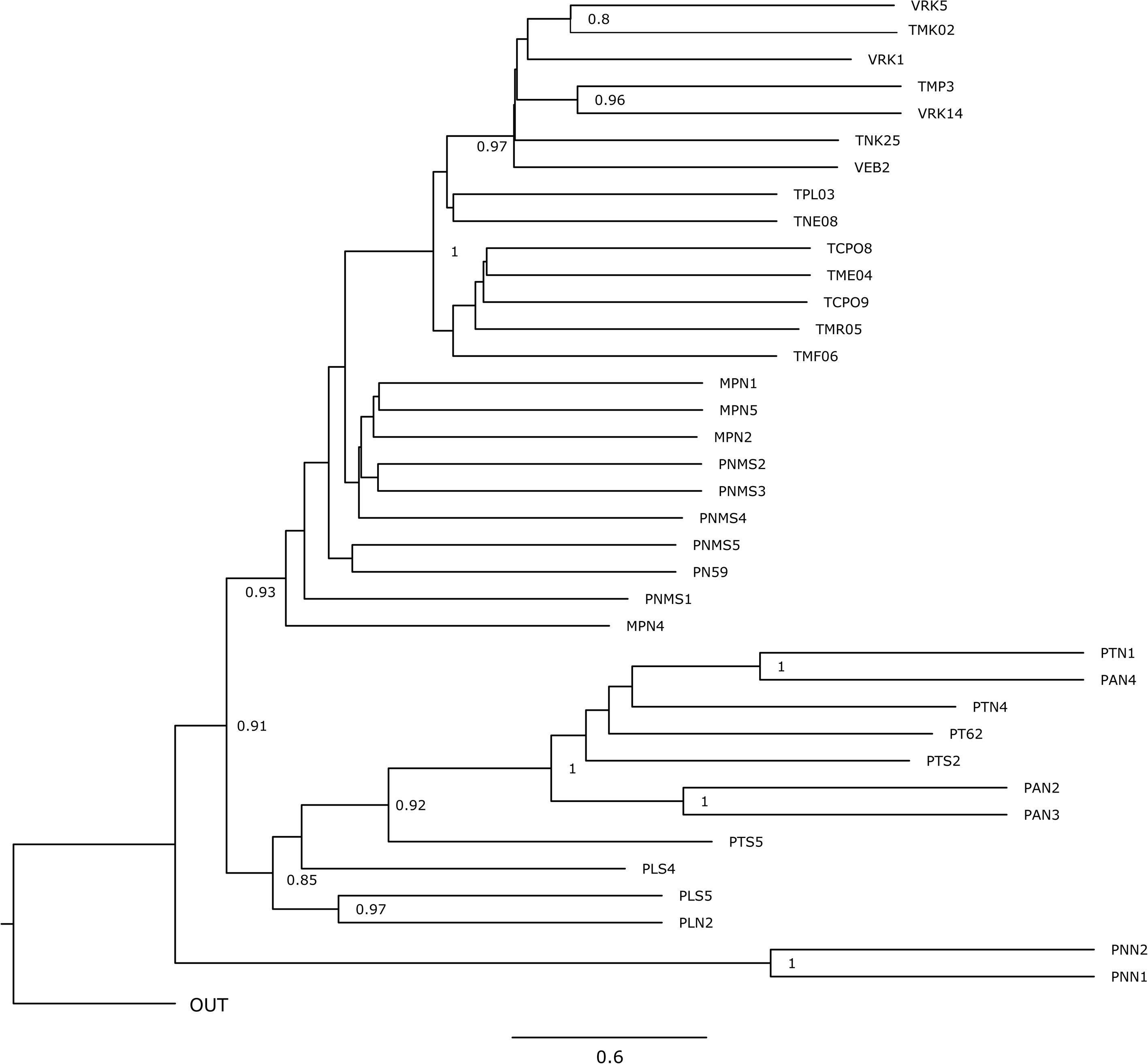
Topology inferred from the nuclear data using the weighted ASTRAL algorithm. Branch lengths are in coalescent units and branch support values (LPP):≥ 80% are shown adjacent to that branch.

**Figure S3:**
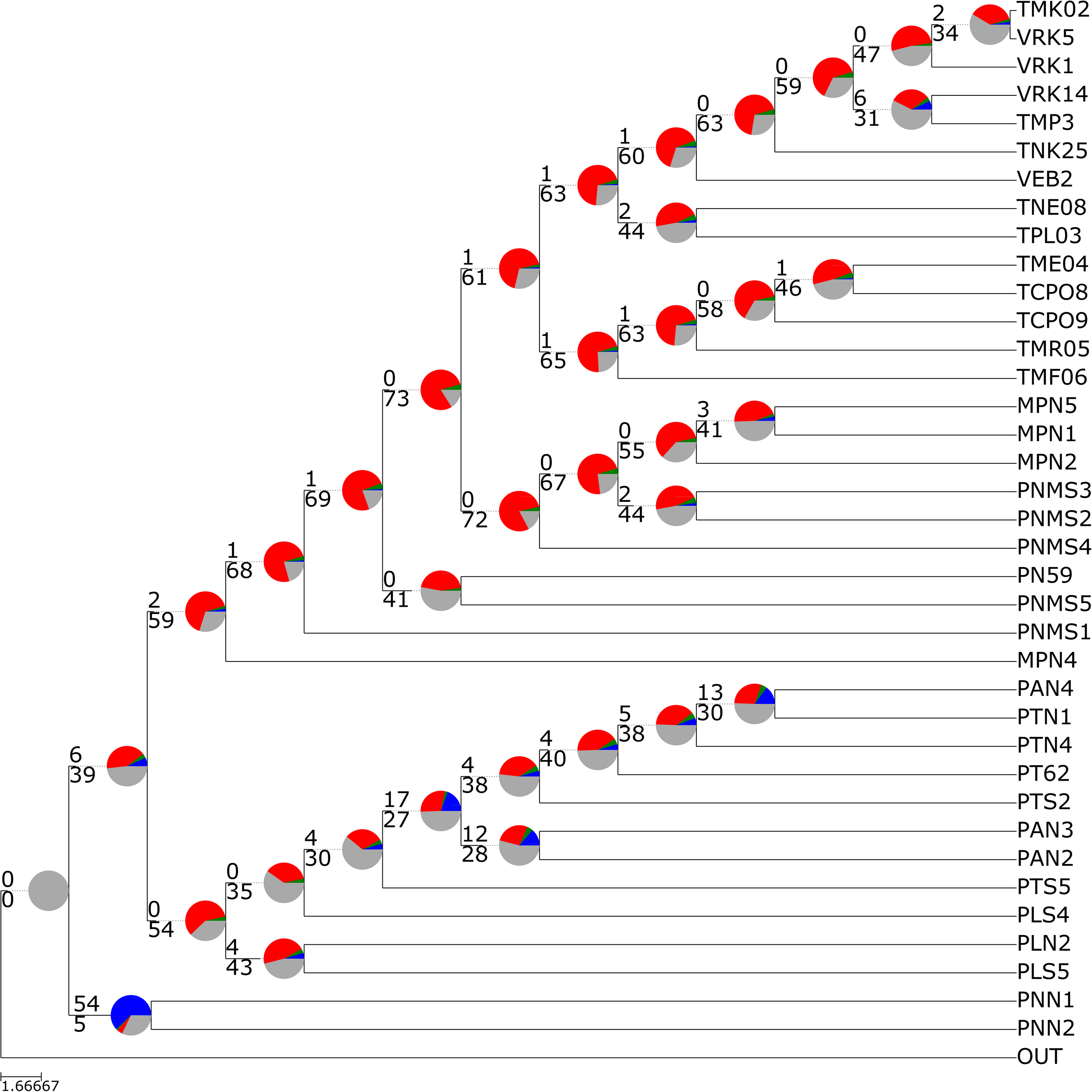
Weighted ASTRAL topology showing the distribution of nuclear gene trees mapped using PhyParts. Numbers of concordant and discordant simulated gene trees are show above and below each branch, respectively. The pie charts represent the proportion of concordant (blue), discordant (main alternative, green; all other alternatives, red) and uninformative (grey) gene tree topologies for that branch.

**Figure S4:**
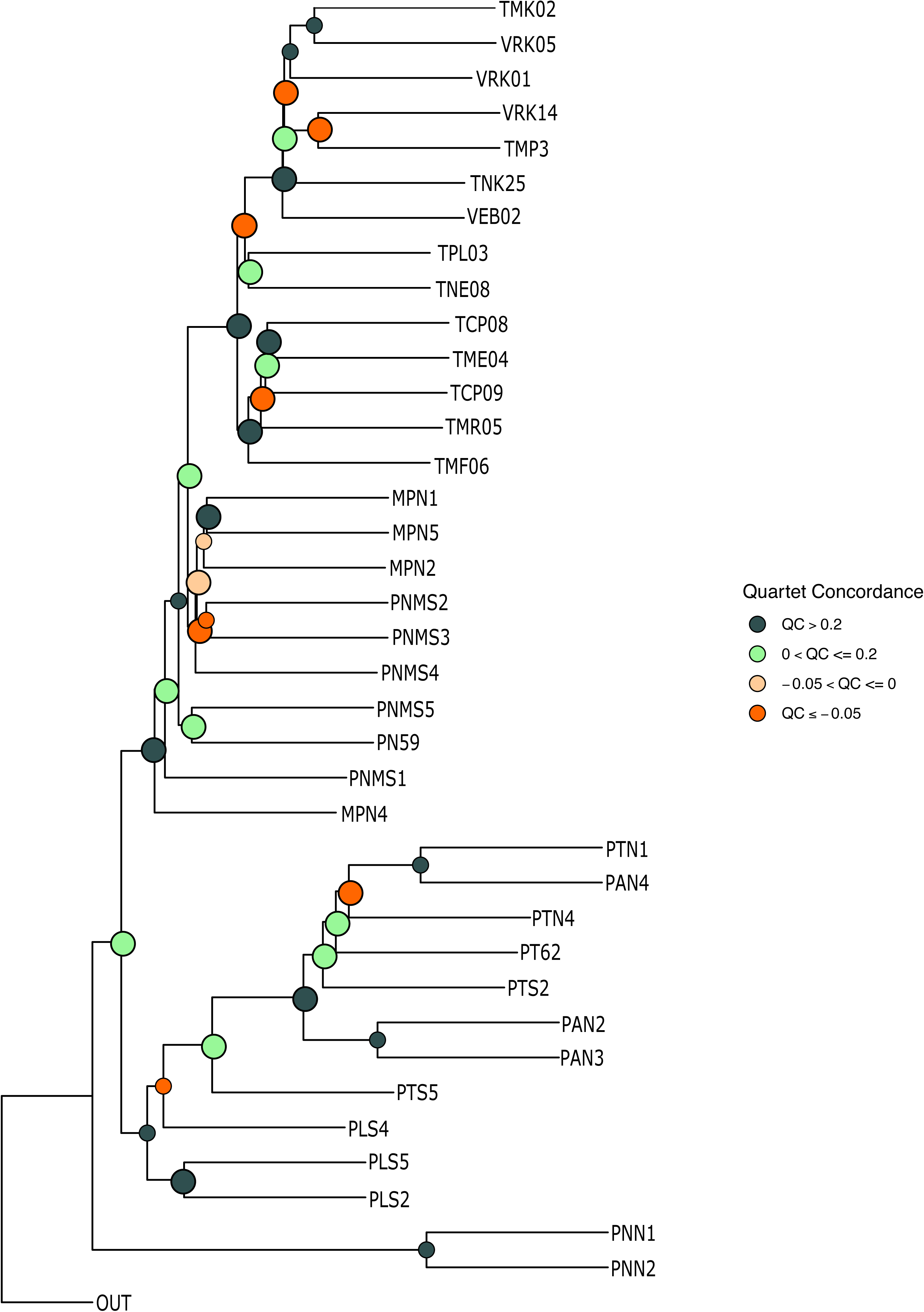
Weighted ASTRAL topology showing heatmap colouration of internal branches for Quartet Concordance (QC). Large node markers had a significant skew according to the *calcQD* test (χ^2^ ≤ 0.05).

**Figure S5:**
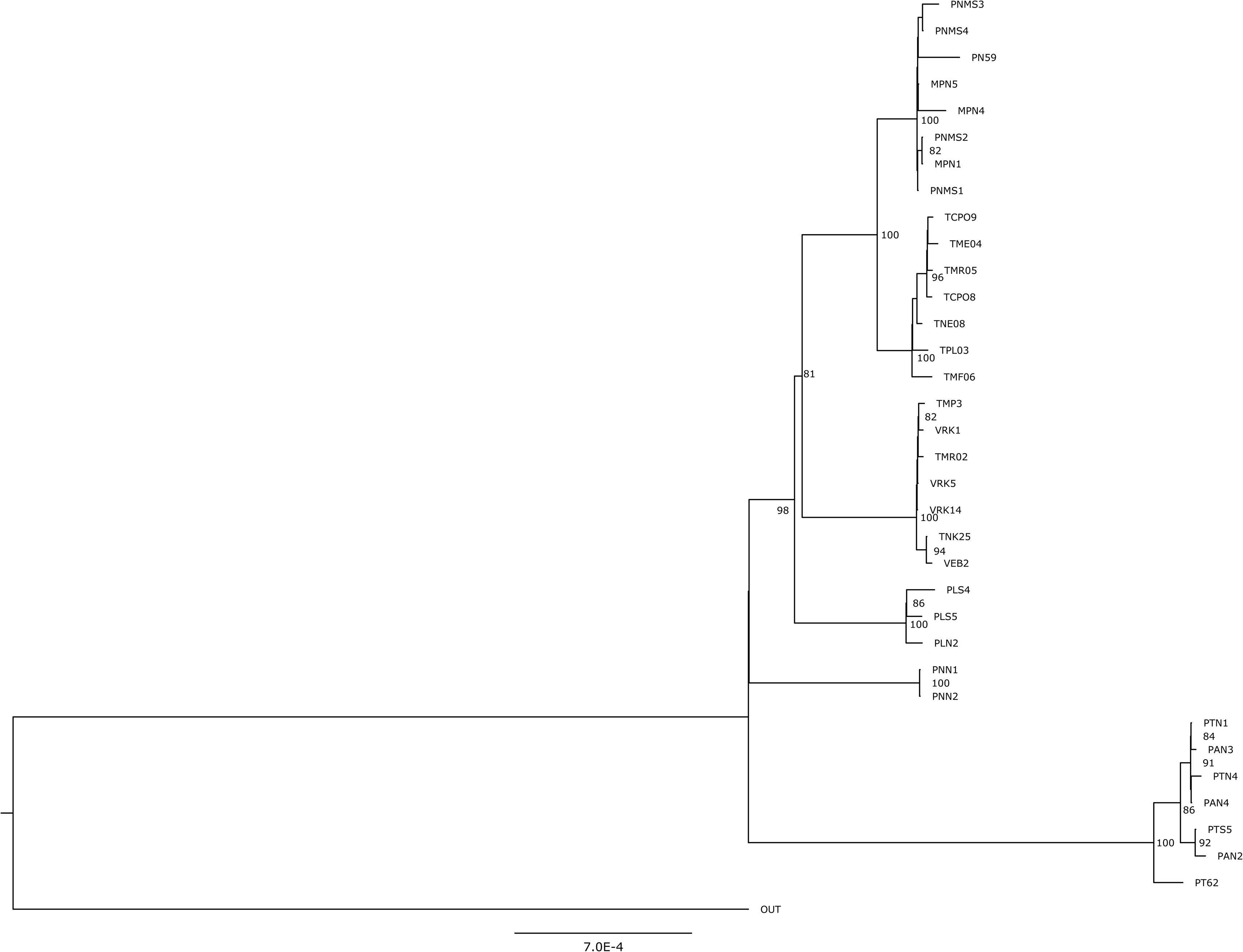
Maximum likelihood topology inferred from the concatenated plastid dataset using IQ-TREE. Branch lengths are proportional to the number of changes occurring on that branch and support values (UFBS):≥ 80% are shown adjacent to that branch.

**Figure S6:**
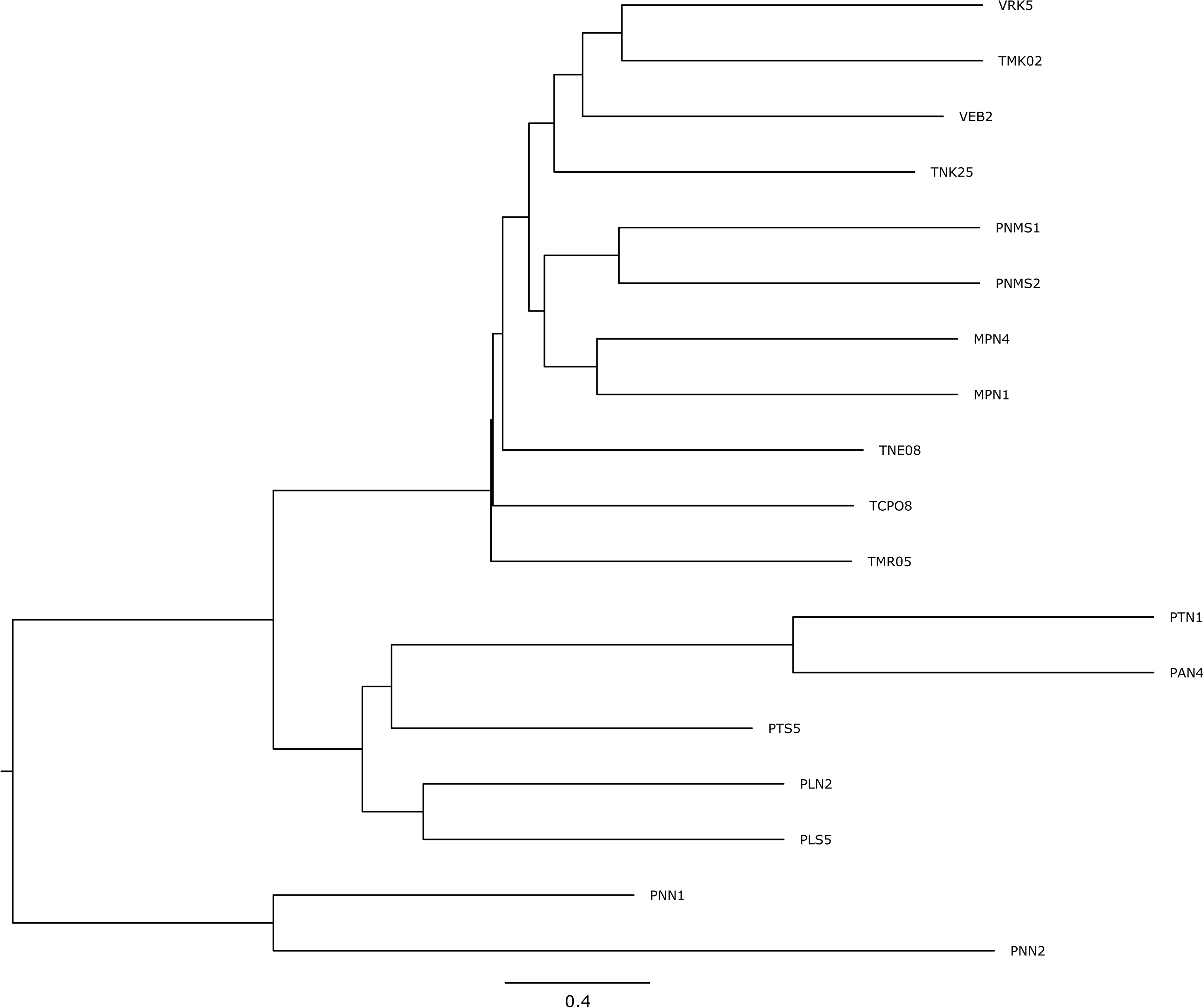
Major tree topology for the best fitting network with two reticulations (*hmax*=2, see Figure 3a) obtained by deleting minor hybrid edges.

**Supplementary Data 1**: Alignment and partition file for the nuclear data set.

**Supplementary Data 2**: Alignment and partition file for the plastid data set.

**Supplementary Data 3**: Weighted ASTRAL tree (‘wASTRAL_bl_4X.tree’) with branch lengths scaled by a factor of 4, 1000 gene trees simulated under a coalescent model using the wASTRAL tree as a guide and the empirical plastid topology.

